# Emissions of climate-altering species from open vegetation fires in the Mediterranean region - A review on methods and data

**DOI:** 10.1101/2024.07.26.605246

**Authors:** Rabia Ali Hundal, Saurabh Annadate, Rita Cesari, Alessio Collalti, Michela Maione, Paolo Cristofanelli

## Abstract

The climate change over the Mediterranean region poses serious concerns about the role of open vegetation fires in the emissions of climate-altering species. The aim of this work is to review the current methodologies for quantifying the emissions of greenhouse gases and black carbon from open vegetation fires, as well as the data provided by four state-of-the-art inventories of emissions of carbon dioxide (CO_2_), methane (CH_4_), nitrous oxide (N_2_O) and black carbon (BC) in the Mediterranean region for the period 2003 - 2020. A limited number of studies specifically addressed the quantification of emissions from open fires in the Mediterranean region. Our data review of fire emissions in the Mediterranean region, where “top-down” methods have not yet implemented, reveals discrepancies across the four inventories examined (GFED v4.1s, GFAS v1.2, FINN v2.5, and EDGAR v8.0). Among these, FINN v2.5 consistently reported the highest emissions, while GFED v4.1s reported the lowest. We observed that the relative ranking of total emissions between the inventories varied for the species considered (e.g. CO_2_ vs. CH_4_) and that different proportions of emissions were attributed to the individual countries included in the Mediterranean domain. We argued that these differences were related to the different spatial resolutions of the input data used to detect the occurrence of fires, the different approaches to calculating the amount of fuel available, and the emission factors used.

The three inventories reporting wildfire emissions were consistent in identifying the occurrence of peaks in the emissions for the years 2007, 2012 and 2017. We hypothesized that La Niña events could partially contribute to triggering the occurrence of these emission peaks.

To increase the accuracy and consistency of climate-altering emission data related to open vegetation fires in the Mediterranean region, we recommend to integrate bottom-up approaches with top-down inversion methods based on satellite and in-situ atmospheric observations.

## 1. Introduction

Since the 19^th^ century, anthropogenic activities have led to a continuous upward trend of greenhouse gas (GHG) emissions (Roelfsema et al., 2020), affecting various aspects of climate, including air and ocean temperatures, sea levels, extreme events related to heat waves and precipitation. In 2022, global anthropogenic GHG emissions reached a new high of 53.8 Gt of CO_2_ equivalent, according to the latest version of the Emissions Database for Global Atmospheric Research (EDGAR v8.0, see Crippa et al., 2023). In 2015, the Paris Agreement established emission targets to keep the rise in global temperature within 1.5°C above pre-industrial levels (UNFCCC, 2015).

According to the latest World Meteorological Organization GHG bulletin (World Meteorological Organization, 2023), the three main GHGs, carbon dioxide (CO_2_), methane (CH_4_) and nitrous oxide (N_2_O), have increased by 150%, 264% and 124% respectively in 2022 compared to 1750. Biomass burning related to open vegetation fires is a relevant source for all of them (Crutzen and Andreae, 1990; Shi et al., 2021). As suggested by Landry (2016), it should be noted that changes in the ecosystem dynamics due to climate change may add further uncertainty to the impact of wildfires on the decadal carbon cycle.

Open vegetation fires also emit dangerous pollutants such as particulate matter, carbon monoxide (CO), nitrogen oxides (NO_x_), sulfur dioxide (SO^2^), and volatile organic compounds, which can harm human health and ecosystems (Anav et al., 2024; Bowman and Johnston, 2005; De Marco et al., 2022). Particulate matter is also associated with major climatic impacts via aerosol-radiation and aerosol-cloud interactions (Intergovernmental Panel On Climate Change, 2023). Globally, biomass burning contributes nearly 30-40% of the atmospheric black carbon (BC), a component of atmospheric aerosol particles resulting from the incomplete combustion of fossil fuels, wood (Bond et al., 2013). By absorbing solar radiation, BC, impacts the climate by heating the atmosphere, accelerating ice melt, and altering cloud properties (Kang et al., 2019, 2020; Klimont et al., 2017). According to Hamilton et al. (2018), reassessment of pre-industrial fire emissions has been shown to have a significant impact on anthropogenic aerosol forcing.

In Europe, emissions from open vegetation fires have been linked to climate variability and change, with anthropogenic influences being an important component (Fernandes et al., 2016; Song et al., 2009). Increasing aridity has been observed to push Mediterranean forest post-fire recovery towards open shrublands (Baudena et al., 2020), aggravating the frequency and intensity of fires in this region as predicted by non-stationary climate-fire models due to global warming (Turco et al., 2019). Wildfires and prescribed burns (controlled fires used for forest management, see Henderson et al. (2005)), the two main types of open vegetation fires that occur most frequently in Europe, both contribute to emissions of pollutants and GHGs. Wildfire intensity is driven by climate change, fuel accumulation, expanding human settlements, flammable vegetation, and prolonged burning (Bowman and Johnston, 2005). The Euro-Mediterranean southern region, where the climate is warmer and drier is where Europe’s wildfires are (and will be) most frequent (Lionello and Scarascia, 2018; Moreira et al., 2011; Noce et al., 2016).

Emissions from open vegetation fires are not as well characterised as other anthropogenic emissions due to the high variability in time and space, the different factors driving activity and the efficiency of emissions (Randerson et al., 2012), including the size and the stage of the fires, the type of vegetation being burned, weather conditions, and human intervention.

Recent studies have highlighted the importance of refining methodologies for emission estimates from open vegetation fires. For instance, advances in satellite-based burned area detection, together with the integration of detailed data on fuel types, meteorological conditions and combustion phases have significantly improved accuracy of bottom-up approaches (Scarpa et al., 2024). Furthermore, innovative observation-informed “top-down” methods incorporating atmospheric chemical transport models have provided new insights into emissions dynamics (Qu et al., 2021, Zhang et al., 2021). Emerging research, such as the study by Pande et al. (2021), demonstrates the effectiveness of integrating satellite data and GIS techniques to estimate biomass resources in semi-arid regions, highlighting the utility for advanced geospatial methods for biomass burning. Despite these advances, significant challenges remain, including biases in emission factors, inconsistencies in detecting smaller fires and limited application of these techniques in the Mediterranean region. These challenges underscore the need for standardised, high-resolution methodologies to improve emission estimates and support climate policies

The different existing methodologies to estimate open vegetation fire emissions can be broadly classified as "bottom-up" or "top-down" (Kasischke and Penner, 2004). The “bottom-up” approach is based on the combination of geophysical (e.g. type of vegetation within a region, proportion of vegetation burned) and biophysical (emission factors) information. The so-called “top-down” approach uses direct atmospheric observations to quantify the net emissions. Traditionally, the “top-down” approach refers to an energy based method. It uses the fire radiative power (FRP) measurements from satellite remote sensing as a direct indicator of the biomass consumed by a fire (Roberts et al., 2005). FRP is then multiplied by a predefined factor, known as the smoke emission coefficient, to estimate pollutant’s emission rates. Rather, in this review, we will refer to the term “top-down” to indicate methods that do not directly use emission factors to calculate emissions, but rather use observations in combination with inverse modeling techniques (Bergamaschi et al., 2018).

To effectively estimate the impact of open vegetation fires on climate, it is essential to have a thorough understanding of the relative performance of different approaches, although each has strengths and limitations. Despite the critical role of fires as a source of GHGs and BC, to our knowledge there is little specific information on open vegetation fire emissions in the Mediterranean region. For this reason, this paper aims to review existing methodologies to quantify GHG and BC emissions from open vegetation fires and data provided by four state-of-the-art emission inventories GFEDv4.1s, GFASv1.2, FINNv1.2 and EDGARv8.0 related to emissions of CO_2_, CH_4_, N_2_O, and BC from open vegetation fires over the Mediterranean region in the period 2003 - 2020.

Although global studies frequently address biomass-burning emissions, a comprehensive regional analysis tailored to the entire Mediterranean region, which is particularly susceptible to climate-driven alterations in fire dynamics, is still unavailable. This paper provides a comparative evaluation of methodologies and data from major emission inventories, focusing on their relative strengths and limitations. By addressing these gaps, this review offers insights to improve emission quantification, particularly for climate altering species, and supports future research and policy making in the Mediterranean region

The paper is structured as follows: in Section 2, we review the methods used to calculate the fire emissions, including both bottom-up and observation-informed “top-down” (inverse modelling) methods, and report on use cases where available.. In Section 3, we present the data review about emissions of climate-altering species in the Mediterranean region related to open vegetation fires. Section 4 discusses the emissions estimates and Section 5 reports our conclusions.

## 2. Methods used for estimating emissions of climate-altering species from vegetation fires: an overview

In this section, we broadly describe two widely used approaches to quantify GHG and BC emissions from open vegetation fires: the so-called “bottom-up” and observation-informed “top-down” approaches.

### 2.1 Bottom-Up

In the bottom-up approach, emissions are derived based on activity data (e.g. area burned, fuel consumption) and emission factors (i.e. a coefficient that describes the rate at which a given activity releases a specific molecule into the atmosphere). This approach is commonly used by environmental scientists and policymakers, offering transparency and replicability in calculating emissions’ contribution for specified areas.

A “bottom-up” system for quantifying open vegetation fire emissions consists of several interrelated modules (see Figure SM1 in the Supplementary Material), each of which provides specific information for a comprehensive emission estimate (van der Werf et al., 2017). These modules include the detection of burned areas using satellite data such as the Visible Infrared Imaging Radiometer Suite (VIIRS) (a sensor onboard the polar-orbiting Suomi National Polar Orbiting Partnership; Suomi NPP, NOAA-21 and NOAA-21 weather satellites) or the Moderate Resolution Imaging Spectroradiometer (MODIS) (a key instrument on board the Terra and Aqua satellites), the calculation of fuel consumption based on biomass burned per unit area and the implementation of emission factors to convert biomass into specific trace gas and aerosol emissions. Temporal distribution modules consistently distribute monthly emissions into daily and hourly values, taking into consideration fire spread rates and diurnal trends. Module outputs are then aggregated into a geospatial framework that maps emissions across landscapes. These components interact through data integration and module calibration to ensure consistency with empirical data and remote sensing. The system provides a comprehensive understanding of the wildfire impacts, including spatially explicit and temporally resolved emissions estimates. By incorporating burned area, fuel consumption, combustion completeness, emission variables, and temporal distribution, the “bottom-up” system improves our ability to monitor and manage the environmental and air quality impacts of wildfires. Volkova et al. (2019) enhanced Australia’s GHG emissions reporting from forest fires using a Tier 2 approach from the 2006 Intergovernmental Panel on Climate Change (IPCC) Guidelines. They refined the emission estimate equations and incorporated detailed, country-specific data from literature reviews and field measurements. This included improvements in parameters such as the area burned, fuel mass, combustion factors, and emission factors. Additionally, they transitioned to a more accurate model for dead organic matter recovery. These refinements aimed to provide precise GHG emission estimates.

A few studies have used a bottom-up approach to estimate GHG, including CO_2,_ CH_4,_ and N_2_O, emissions from open vegetation fires in Europe.

A study by Rosa et al. (2011), estimated atmospheric emissions from wildfires in Portugal from 1990 to 2000. The researchers used Landsat-based burnt area maps, land cover maps, national forest inventory data, biometric models, and emissions factors from literature. Emissions were calculated based on the area burnt, biomass loading, combustion factor, and emission factor specific to land cover types. The results showed a strong correlation between the area burnt and the emissions, with the combustion factor for shrubs and emission factors identified as the primary sources of uncertainty compared to other sectors, wildfire emissions constitute 1% to 9% of Portugal’s total annual GHG emissions.

Scarpa et al. (2024) employed a bottom-up approach to estimate GHG and particulate matter emissions from rural and forest fires in Italy between 2007 and 2017. Their method improved emission accuracy by integrating detailed data on burned areas, fuel types, meteorological conditions and combustion phases, offering results that aligned well with global and national inventories. According to this study, fire disturbance in broadleaf forests, shrublands, and agricultural fuel types is the main source of vegetation fire emissions in Italy, accounting for about 76 % of the total. They estimated average annual emissions from open vegetation fire by 2346 Gg year^-1^ for CO_2_ and 10.4 Gg year^-1^ for CH_4_.

### 2.2 Top-Down

The observation-informed "top-down" approach is a method in which atmospheric observations are used to improve the emission estimates of the atmospheric species of interest. This approach estimates emissions by adopting a framework that integrates atmospheric observations, an atmospheric chemical transport model, and an inversion algorithm. The approach is based on the relationship between atmospheric concentrations or mole fractions with emissions, i.e. the so-called ‘source-receptor relationships’ calculated by the chemical transport model. The top-down method uses a “bottom-up” inventory as prior knowledge and optimises it to match the modelled concentrations with observations. By leveraging the combined information from both “bottom-up” estimates and observations, the inverse modelling approach significantly reduces uncertainties in the posterior emission estimates (Rodgers, 2000). The accuracy of these inverse modelling results and the spatial scale at which emission estimates are obtained depend on the quality and spatial density of the measurements and the precision of the transport model used in the analysis (Bergamaschi et al., 2018).

Modelled mixing ratios corresponding to the observations can be calculated as:

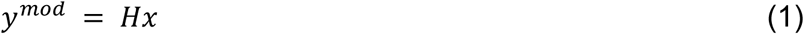

where *y^mod^* is a vector of the modelled mixing ratio, *x* is the unknown emission flux vector and H is the source receptor relationship matrix. The emissions fluxes vector *x* includes the variables to be optimised by the inversion, i.e. gridded emissions and other elements such as background mixing ratio values.

Observed values can be expressed as:

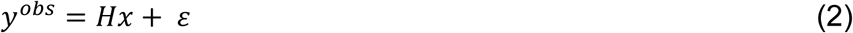

where *y^obs^* is the observation vector and *ε* represents the observation error. However, observations might not be sufficient to precisely constrain all components of the emission vector. This situation leads to an ill-conditioned problem, requiring regularisation or the introduction of supplementary information to obtain a meaningful solution. Typically, this additional constraint is provided as a priori estimates of the unknowns, i.e. spatially distributed information of the fluxes (*x_b_*), obtained from the bottom-up emission estimate. Following Bayes’ theorem, the most likely solution of the posterior emission flux (*x*) minimises the difference between the observed and modelled mixing ratios. Assuming that the observation and prior uncertainties follow Gaussian probability density functions, this can be formulated as a cost function:

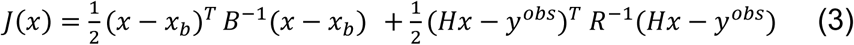

In this equation, *B* represents the prior flux error covariance matrix, and *R* is the observation error covariance matrix. Several methods can be employed to find the optimum posterior state, defined as the value of *x* for which eq. 3 has a minimum. Common approaches include the analytical inversion (Stohl et al., 2009), Kalman filter (Dash et al., 2024), 3D-Var (Three-Dimensional Variational Assimilation) and 4D-Var (Four-Dimensional Variational Assimilation) (Meirink et al., 2006) optimisation techniques.

Inversion models have evolved rapidly in recent years, reducing uncertainties in emission estimates. For a long time, the inversion methods were used to optimise the net emissions of the species of interest. However, there have been advancements in the field to accommodate the extraction of the sectorial contributions to the posterior total emissions. In a commonly used and straightforward approach, net fluxes are partitioned to underlying sector-based emissions by scaling fluxes based on the relative weight of sectors in a prior inventory (Qu et al., 2021). This does not assume that the prior distribution of sectoral emissions is correct, only that the relative allocation within a given grid cell is correct. A similar approach accounts for emissions from different sectors having different prior error standard deviations contributing differently to each sector (Shen et al., 2023). In another approach by Zhang et al. (2021), the state vector is split into different emission sources, such as wetland and non-wetland emissions. It allows the emission sector with higher uncertainty to be treated differently than the well-known sector. In a similar approach by Lu et al. (2021), the state vector is organised in three ways to obtain monthly, sectorial, and provincial emissions according to the subnational boundaries for each year. Assuming relative weights of different sectors in prior emissions imposes a correlation between emission sectors that may not exist. Cusworth et al. (2021) discussed a method that projects inverse CH_4_ fluxes directly to emission sectors while accounting for uncertainty structure and spatial resolution of prior fluxes and emissions. This method can also be helpful in partitioning highly uncertain biospheric and anthropogenic emissions to reconcile the global budget.

In more sophisticated approaches to get accurate sectorial estimates, the inversion system is generally provided with some additional constraints. For instance, to constrain CH_4_ fluxes, additional trace gases can be employed to partition emissions. This method becomes feasible when these trace gases are co-emitted from a specific source at a characteristic ratio. For example, CO, the classical tracer for tracking smoke plumes from fires, can be used to estimate methane emissions from activities such as biomass burning (Heald et al., 2004). Similarly, ethane (C_2_H_6_) is emitted during fossil fuel production and use, without significant emissions from biogenic sources used to quantify fossil fuel emissions (Helmig et al., 2016; Peischl et al., 2013). Observations of isotopologues can also be harnessed to apportion emissions through a similar approach. This method involves evaluating the ratios of isotopologues emitted from different source types. Lan et al. (2021) used CH_4_ isotopologue measurements to better understand and differentiate emissions from various sources, including biomass burning emissions. Analysing these isotopic signatures provides valuable information on the sources and a more comprehensive understanding of methane emissions. To our knowledge, for the Mediterranean region, no “top-down” quantification based on inversion frameworks exists for the emissions of climate-altering species from the open vegetation fires.

## 3. Quantification of open vegetation fire emissions over the Mediterranean basin: a data review

We present a synthesis of the degree of agreement between different state-of-the-art inventories estimating emissions of climate-altering species (CO_2_, CH_4_, N_2_O and BC) from open burning in the Mediterranean region. To this end, we compared emissions for the whole area as well as for individual countries for four inventories (GFAS, GFED, FINN, and EDGAR) from 2003 to 2020. Table 1 summarises key characteristics of the considered inventories, including their acronyms, versions, producers, coverage details, and resolutions. More specific details about each inventory are provided in the supplementary material (Table SM1). Note that GFED, CAMS, and FINN include anthropogenic and natural vegetation fires, whereas EDGAR only includes anthropogenic emissions related to agricultural practices.

**Table 1.**
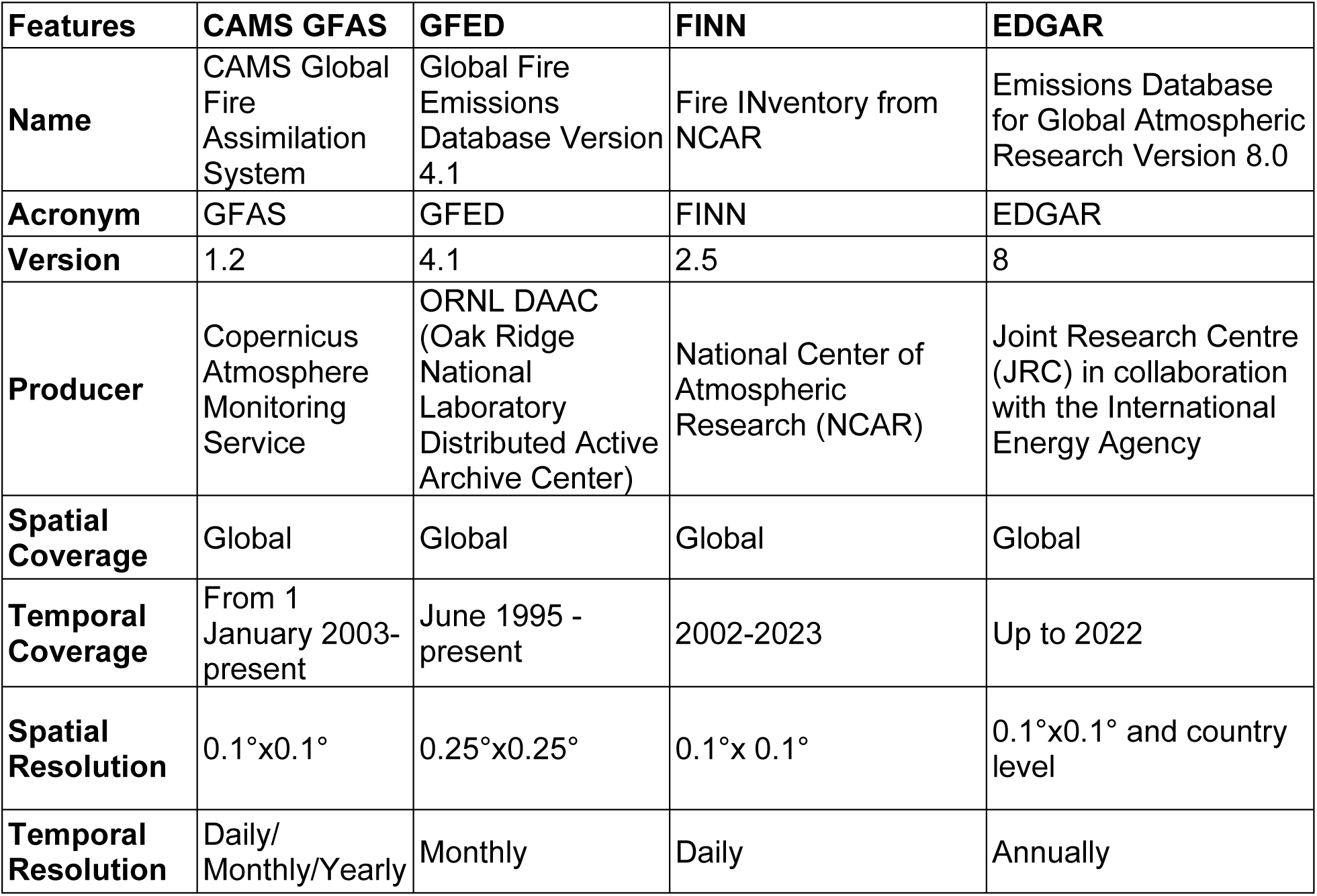
Overview of fire emission inventories considered in the comparison.

The Mediterranean region considered for aggregating emissions is shown in Figure 1. It is based on HydroSHEDS (Hydrological data and maps based on Shuttle Elevation Derivatives at multiple Scales) developed by the World Wildlife Fund (Lehner and Grill, 2013), a global dataset that provides detailed hydrographic information, like river networks and watershed boundaries, in a consistent format for large scale applications. The definition of the Mediterranean region is not unique and different definitions can be adopted depending on the purposes and discipline (Giorgi, 2006; Lefèvre and Fady, 2016; Meybeck et al., 2006; Schicker et al., 2010). In this case, we defined a domain based on the extension of the catchment area of rivers flowing into the Mediterranean Sea, including the lower part of the Nile catchment. This definition is a compromise between other definitions and is consistent with Schicker et al. (2010).

**Figure 1.**
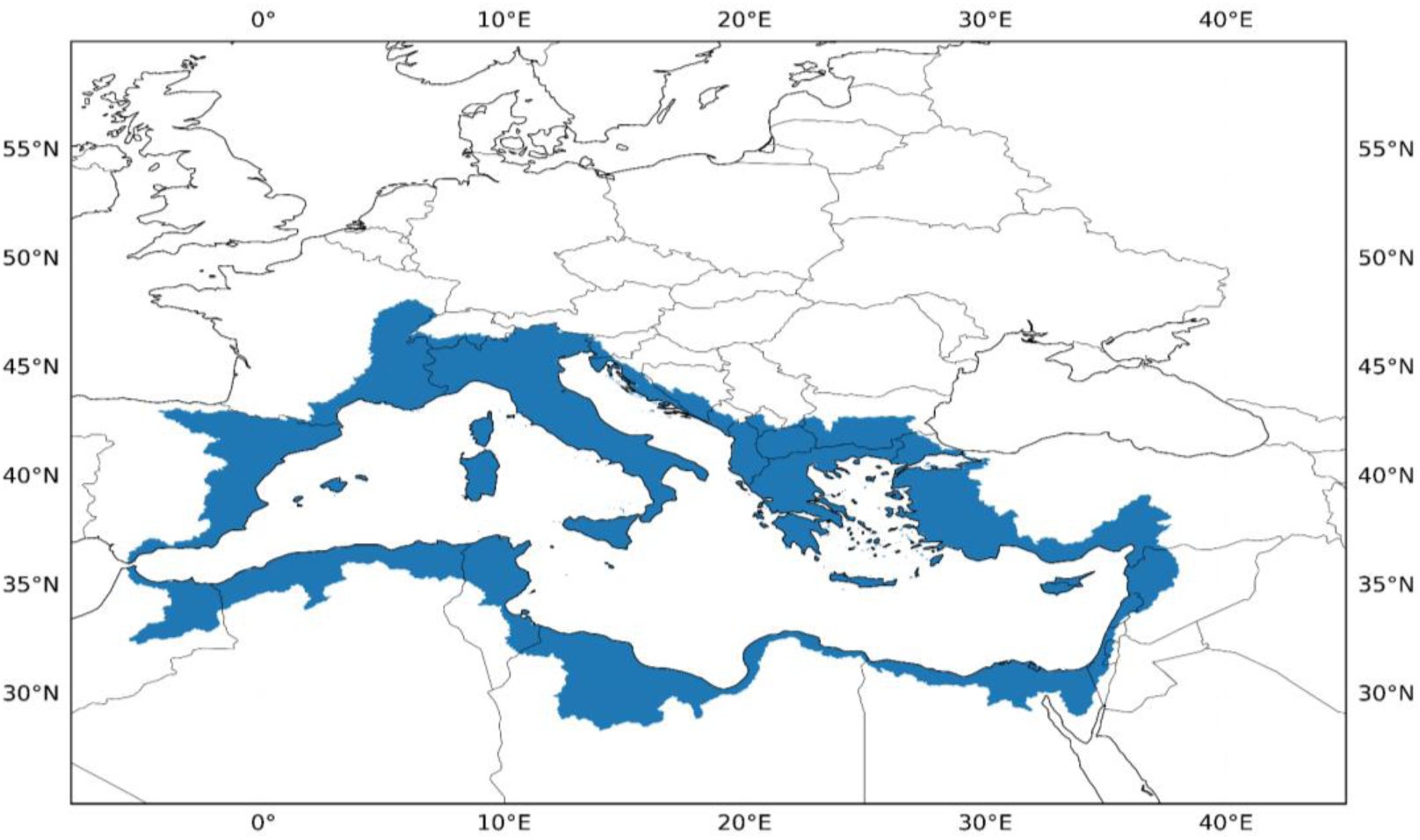
The Mediterranean region domain (blue area in the map) selected to aggregate emission fluxes from CAMS, GFED, FINN and EDGAR emission inventories.

The area of the Mediterranean that we have considered represents about 1.5% of the global land surface. Using global CO_2_ emissions from wildfires provided by Samborska et al. (2024) as a reference, the percentage contribution of Mediterranean wildfires to global wildfires emissions during the study period ranged from a minimum value of 0.18 - 0.32% in 2012 to a maximum value of 0.24 - 0.45% in 2007, depending on the inventory considered. It should be noted that these numbers refer to the domain as defined above and may change depending on the spatial domain selected to represent the Mediterranean region. For example, the relative values increase by including the whole Iberian peninsula in the analysis as done by Gudmundsson et al. (2014) and Turco et al. (2018).

Figure 2 reports the multi-annual variability of open fire emissions for CO_2_, CH_4_, N_2_O, and BC over the Mediterranean region from 2003 to 2020.

**Figure 2.**
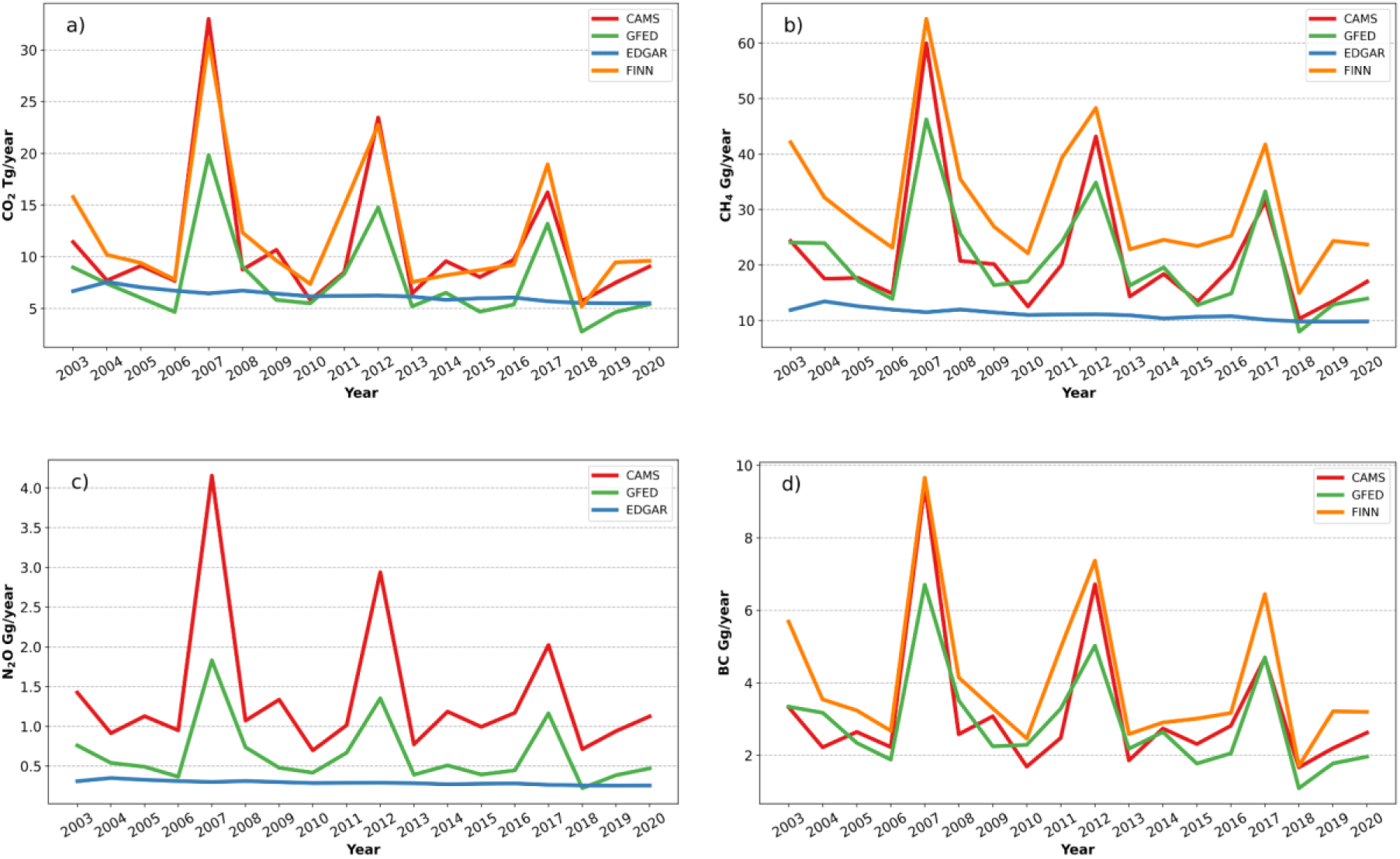
Annual emissions from open vegetation fires in the Mediterranean region (2003-2020): CO_2_ (a), CH_4_ (b), N_2_O (c) and BC (d). The different lines denote CAMS (red), GFED (green), EDGAR (blue), and FINN (orange).

The GFED, CAMS, and FINN inventories are consistent in identifying temporal peaks in the emissions, although there are differences in absolute values, with CAMS and FINN often showing emission levels higher than GFED. EDGAR showed higher CO_2_ emission values than GFED for 10 out of 18 years. This is surprising, as EDGAR is expected to represent a lower limit for emissions, as it only reports agriculture vegetation fires. EDGAR does not capture the significant emission peaks observed in 2007, 2012 and 2017, probably because it only includes emissions related to agriculture and waste burning.

In 2007, “extreme” fire danger conditions affected south-eastern Europe at the end of July and August (Cesari et al., 2014). These conditions led to catastrophic events affecting southern Italy and Greece, with record numbers of burnt areas. In 2007, the Rapid Damage Assessment module of the European Forest Fires Information System reported high burnt areas for Greece (271,516 ha) and Italy (153,753 ha) (European Commission et al., 2008). The weather conditions that contributed to the wildfires included prolonged heat waves with very high temperatures, prolonged droughts that preceded the events, and strong winds that coincided with the ignition of the fire (Koutsias et al., 2012).

According to Schmuck (2013), in 2012 forest fires in the five southern EU countries (Portugal, Spain, France, Italy, and Greece) burned a total area of 519,424 ha. This value was well above the average for the previous 20 years (∼ 400,000 ha) and among the highest since 2000 (only 2003, 2005, and 2007 reported higher values of burnt areas, see San-Miguel-Ayanz et al. (2022)). The winter and early spring were marked by a severe drought in France, Spain and Italy with the period June to August hotter and drier than normal (Schmuck, 2013).

In 2017, as reported by the European Drought Observatory, Italy experienced a severe drought, followed by one of the driest springs in the last 60 years. During this period, some regions received 80% less rainfall than normal. Two-thirds of Italy was affected by these conditions (Joint Research Center, 2024) leading to a burnt area of 140,404 ha with 788 fires (Libertà et al., 2018). Interestingly, while 2007, 2012 and 2017 were years with significant peaks in burned areas, the trend across these years shows a decrease in the magnitude of these peaks. This decreasing trend is consistent with the behaviour observed for emissions of key climate-altering species including CO_2_, CH_4_, N_2_O and BC.

Figure 3 shows the percentage of contributions of different countries to the CO_2_ emissions from the Mediterranean region over the period 2003 - 2020. CO_2_ has been presented as a ’representative’ case because it is the dominant GHG emitted by open burning and is a major contributor to global warming and climate change. Results for CH_4_, N_2_O and BC can be found in the supplementary material (Fig. SM2). It should be noted that EDGAR was not included in this figure, as it only provides emission estimates for agricultural waste burning and does not account for emissions from other large fires. After extracting the Mediterranean region from each global inventory, we segregated national emissions falling in the domain defined by Figure 1.

**Figure 3.**
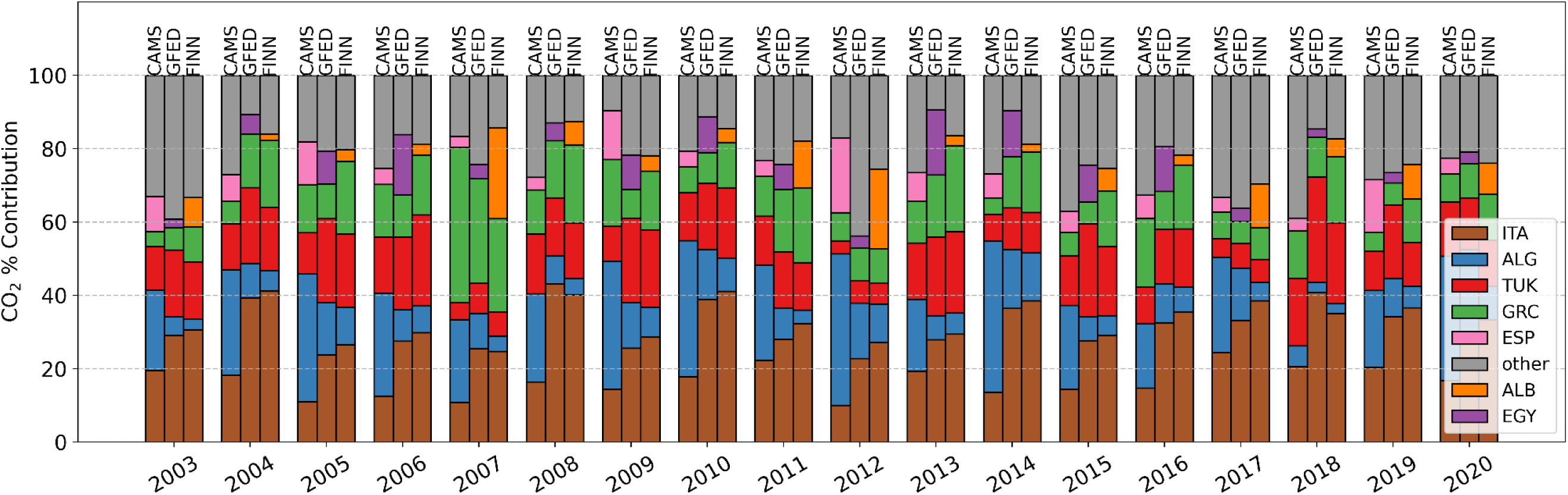
Annual CO_2_ emissions contribution for the top 5 emitting countries for CAMS/GFAS v1.2 (left column), GFED v4.1s(middle column) and FINN v2.5 (right column) for 2003 - 2020 in the Mediterranean region (Albania: ALB, Algeria: ALG; Egypt: EGY; France: FRA; Greece: GRE; Italy: ITA; Spain: ESP, Turkey: TUR; other countries: OTH).

An interesting question is the degree of agreement between the different inventories when looking at individual countries included in the Mediterranean area. Here we discussed the case for CO_2_ emissions (Figure 3) as a representative example. Over the whole investigation period, CAMS identified Algeria as the most important contributor for CO_2_, while both GFED and FINN indicated Italy. However, differences were found for the different years. As an example, in 2012 which is characterised by the occurrence of a peak in emissions, all inventories indicated Italy as a prominent contributor, but with varying proportions: CAMS reported 24%, GFED 33% and FINN 39% of the total emissions. A notable difference is that CAMS reported Spain as one of the top five Mediterranean contributors for each year, while GFED reported Egypt and FINN reported Albania. These discrepancies highlight the challenges of reconciling data from different inventories and the need for further scrutiny of the methodologies used by each inventory especially when looking at smaller spatial scales. It should be emphasised that the aim of this country-level analysis is not to rank the contributors to emissions from open burning of vegetation (this should take into account the different areas of the countries), but to highlight the discrepancies in the estimates provided by the inventories here reviewed.

As suggested by Magi et al. (2012), through agricultural and waste burning, human practices add further complexity to occurrence of vegetation fires, with human burning practices in agriculture not necessarily following the temporal development of wildfires. As EDGAR provides emission estimates for agricultural field burning and not those from other large fires (e.g. biomass burning from Savannah and forests, see Petrescu et al. (2020)), it will be used in the following to analyse the emissions of CO_2_, CH_4_, and N_2_O from the agriculture sector in the period 2003 - 2020 (Figure 4).

**Figure 4.**
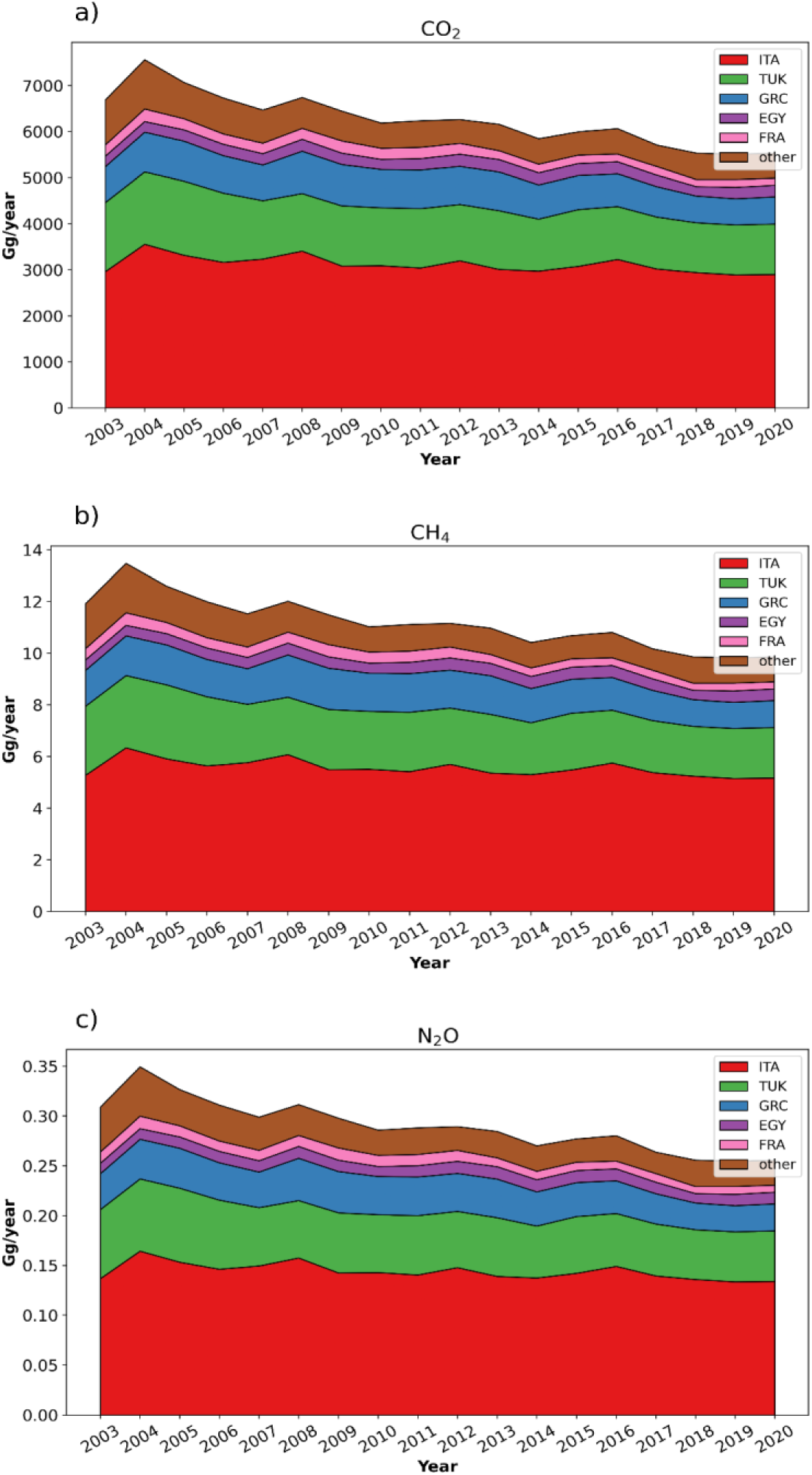
Time series of agriculture and waste burning emissions reported by EDGAR v.8 for different countries.

Table 2 illustrates the trends in emissions of CO_2_, CH_4_ and N_2_O over the period 2003 to 2020 using EDGAR inventory. The trends are estimated using the Theil-Sen slope estimator (Wilcox, 2010) and are presented for individual top 5 emission contributing countries (France, Greece, Italy, Turkey, Egypt) and as an aggregated total for the Mediterranean domain. The results show statistically significant downward tendencies for most countries, with Turkey showing the largest decreases. The aggregated total emissions reveal a reduction in CO_2_ (-0.09 Tg year^-1^), CH_4_ (-0.16 Gg year^-1^) and N_2_O (-4.2 Mg year^-1^) emissions.

**Table 2.** Theil-Sen trend analysis of anthropogenic emissions (2003 - 2020) from the EDGAR inventory for Egypt (“EGY”), France (“FRA”), Greece (“GRC”), Italy (“ITA”), Turkey (“TUK”), and the whole study domain (“MED”). Asterisks (*, **, ***) represent statistical significance in confidence levels at p<0.05, p<0.01 and p<0.001, respectively. The 95% confidence intervals in the slopes are reported within brackets. Emissions for countries refer to the part of the national territory included in the study area shown in Figure 1.

| Spatial region | CH<sub>4</sub> Trend (Gg yr<sup>-1</sup>) | CO<sub>2</sub> Trend (Tg yr<sup>-1</sup>) | N<sub>2</sub>O Trend (Mg yr<sup>-1</sup>) |
| --- | --- | --- | --- |
| EGY | $1.4 \times 10^{-3}$<br>[-2.1 $\times 10^{-3}$, 4.6 $\times 10^{-3}$] | $7.6 \times 10^{-4}$<br>[-1.2 $\times 10^{-3}$, 2.6 $\times 10^{-3}$] | -0.0 [-0.1, 0.1] |
| FRA | -0.01 [-0.02, -0.01]*** | -0.01 [-0.01, 0]*** | -0.3 [-0.4, -0.2]*** |
| GRC | -0.03 [-0.05, -0.01]** | -0.02 [-0.03, -0.01]** | -0.8 [-1.2, -0.3]** |
| ITA | -0.04 [-0.07, -0.01]* | -0.02 [-0.04, 0]* | -1.0 [-1.9, -0.2]* |
| TUK | -0.05 [-0.06, -0.03]*** | -0.03 [-0.04, -0.02]*** | -1.2 [-1.6, -0.8]*** |
| MED | -0.16 [-0.21, -0.13]*** | -0.09 [-0.21, -0.07]*** | -4.2 [-5.4, -3.3]*** |

This reduction of the emissions from the European countries in the domain is consistent with the implementation of directives defined to control agricultural and waste burning, as highlighted by the United Nations Economic Commission for Europe and supported by the European Union’s sustainable waste management policies (United Nations Environment Programme, 2024).

## 4. Discussion on emission estimates

Vegetation fires are a complex process associated with factors of different origins and occurring on different spatial and temporal scales, such as vegetation and landscape conditions, weather conditions, and human activities (Turco et al., 2013). It is thus difficult to quantify and univocally attribute the relative importance of these different processes to fire occurrences and related emissions. The data comparison reported in Section 3 showed that all three considered inventories that include wildfire contributions (i.e. GFED, GFAS and FINN) showed temporally consistent emission peaks for the years 2007, 2012 and 2017. Scarpa et al. (2024) identified these years as the most significant over the period 2007 - 2017 in terms of GHG and particulate aerosol emissions from vegetation fires over Italy. Previous studies suggested that climatic processes, through their impact on fuel moisture, as well as on nature and availability of fuel, can represent a driving process for the inter-annual variability of regional fire patterns, especially in the Mediterranean region (Meyn et al., 2007). In particular, a ∼5-year recurring maxima in the burned areas was observed by Turco et al. (2019) for the northeastern Iberian Peninsula related to summer precipitation and maximum temperature as well as antecedent climate conditions with a time lag of 1-2 years. From this perspective, it is interesting to note that all the years that are characterised by a peak in open vegetation fire emissions over the Mediterranean basin are featured by the occurrence of La Niña events. La Niña represents the “cold” phase of the El Niño Southern Oscillation (ENSO), a quasi-periodic (two to seven years) climate pattern affecting the equatorial Pacific circulation (Burton et al., 2020). According to NOAA, the magnitude of ENSO events is classified into categories such as "weak", "moderate", "strong", or "very strong". Based on this classification, the 2007 - 2008 event was categorized as “strong” ENSO, the 2011 - 2012 event as “moderate”, and the 2017 - 2018 event as “weak” (Climate Prediction Center, 2024). Particularly cold phases of ENSO (i.e. La Niña) have occurred in May 2007 - May 2008, June 2011- May 2012 and June - December 2016 (Climate Prediction Center, 2024), i.e. at the same time as or in advance (with a lag time up to 1 year) of the observed peak fire seasons (Figure SM3 in the supplementary material).

ENSO is the primary cause of fluctuations in the terrestrial carbon cycle, particularly in the tropics (Cox et al., 2013): previous studies (Le Page et al., 2008; Zhai et al., 2016) suggested a significant global influence of ENSO on the carbon budget by increased fire activity and decreased terrestrial carbon sink capacity.

For the Asia-Pacific region, ENSO has been an important climate driver, but its influence over North Atlantic Europe is quite challenging to establish (Ashok et al., 2007; Ayarzagüena et al., 2018), despite previous research suggesting a robust climate influence over Europe (Brönnimann, 2007).

The South Asian summer monsoon and ENSO are closely related through atmospheric teleconnections. ENSO modulates the strength and intensity of monsoons with la Niña events leading to stronger monsoons. This variability in the monsoon, in return, impacts the Mediterranean climate through the monsoon-desert mechanism, affecting subsidence and precipitation patterns in the region. In the years with a stronger monsoon, there is a noticeable increase in the vertical advection of moist static energy and horizontal advection of dry enthalpy over the Mediterranean region, leading to drier conditions (Cherchi et al., 2014). Shaman (2014), showed decreased precipitation during La Niña events over southern Mediterranean Europe in June - September and October - December.

Thus, despite great caution should be exercised when discussing possible climatic processes over the relatively short study period considered in this study, we argue that ENSO may have played a role in determining part of the observed inter-annual variability of GHG and BC emissions associated with open vegetation fires in the Mediterranean region. With the aim of providing, also in the context of this review, indications to better disentangle the possible relationship between ENSO and regional climate variability with wildfire emissions, we calculated the anomalies of surface air temperature (2m temperature) and total precipitation for the Mediterranean area. For this analysis we utilised data from the ECMWF fifth generation reanalysis, i.e. ERA5 (Hersbach et al., 2020), with a spatial resolution of 0.25° x 0.25°. The mean of ensemble members from the dataset “ERA5 monthly averaged data on single levels from 1940 to present” (Hersbach et al., 2023) provided by the Copernicus Climate Change Service was used. Anomalies were calculated for the season May - August with respect to the means in the referenced period. Years 2012 and 2017 were characterised by evident positive and negative anomalies for temperature and precipitation, respectively (Fig. 5). No clear anomaly signals were observed for 2007. These results did not change dramatically according to the length of the seasons analysed (see Fig. SM4, in the Supplementary Material). It should be noted that the 2007 fire season was mainly influenced by specific large events that occurred over Italy and Greece in July and August (Cesari et al., 2014; European Commission et al., 2008; Koutsias et al., 2012). It is therefore possible that our Mediterranean-scale anomaly may not fully represent the weather variability occurring at the sub-domain and sub-seasonal scales. Furthermore, it should be considered that, as suggested by Turco et al. (2013), several other factors besides current weather conditions may influence the occurrence of wildfires in the Mediterranean, including weather conditions preceding (up to 2 years before) the actual fire season, as well as human management, including fire suppression activities, which may affect fuel availability and type/conditions. Finally, Zhang et al. (2019), pointed to the contrasting responses of La Niña events in influencing the extratropical atmospheric circulation, adding another layer of complexity in establishing a cause-and-effect relationship.

**Figure 5.**
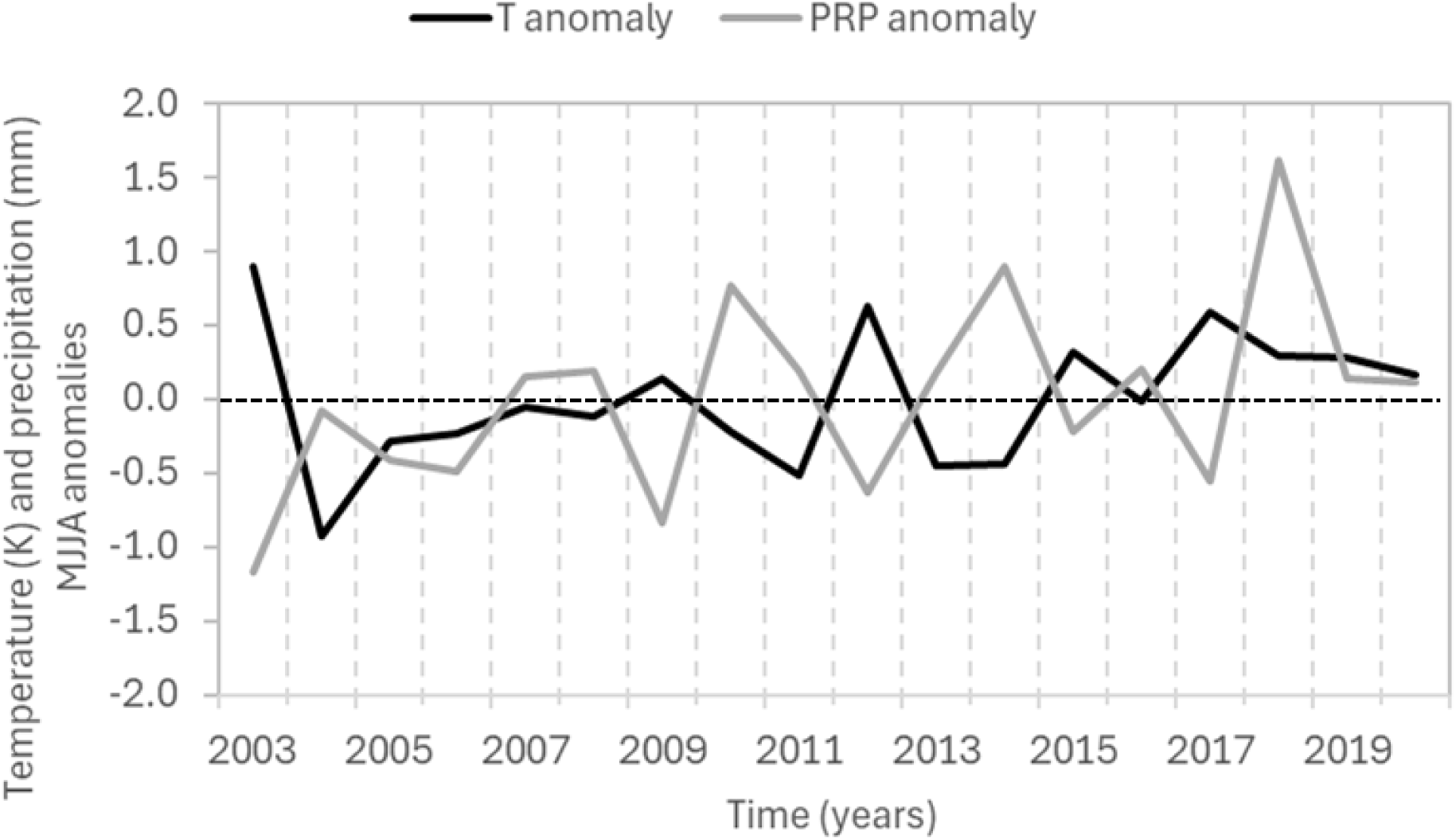
Surface temperature and precipitation anomalies from May to August over the period 2003 - 2020 for the Mediterranean domain as calculated from the ERA5 dataset.

It should be noted that our integrated basin perspective and the annual temporal resolution adopted to compare the emission inventory in this review paper, prevented us from considering specific emission variability that can occur among different source regions and on sub-seasonal time scales (Gudmundsson et al., 2014).

Despite their agreement on the diagnosis of peak years in emissions, there were notable differences in the quantification of total GHG and BC emissions in the Mediterranean for GFED, CAMS, and FINN. On average, FINN provides the highest emission estimates for all species considered, while GFED provides the lowest, even if it includes emissions from small fires. For CO_2_ (CH_4_), more consistent emission estimates are provided by FINN (GFED) and CAMS. The differences in GHG and BC emission among GFED, CAMS GFAS, and FINN might originate from variations in methodologies and input data sources. GFED estimates burned area and emissions using satellite imagery, with a spatial resolution of 0.25° x 0.25°, covering a range of fire types and vegetation. CAMS relies on fire radiative power data from MODIS instruments to estimate emissions at a finer resolution of 0.1° x 0,1°. FINN provides emissions estimates with high spatial resolution (1 km^2^), integrating fire detections from MODIS and Visible Infrared Imaging Radiometer Suite (VIIRS). Thus, it can be argued that the higher spatial resolution of satellite data used to obtain the burned area could partially explain the higher values of emissions. However, it should be considered that each dataset applies different methods to calculate the fuel amount as well as emission factors tailored to vegetation types and fire characteristics. In particular, the differences in the emission factors can explain the different relative agreement between the reviewed inventories when different species are considered (e.g. CO_2_ vs CH_4_).

To assess the spread of the four emission inventories, we calculated the mean standard deviation (σ) and the mean normalized standard deviation (coefficient of variation, v) for the study period for the top three emitting countries and the entire study domain (Table 3). The σ parameter provides indication about the temporal variability of emissions during the investigation period. The coefficient of variation (v) can be used as a measure of the emission spread across the different inventories for the species.

**Table 3.** Average standard deviation (σ) and average normalised standard deviation (coefficient of variation, v) for Algeria (“ALG”), Greece (“GRC”), Italy (“ITA”), and the whole study domain (“MED”) for different atmospheric species. σ is in Tg for CO_2_ and in Gg for all other species.

| Species | ALG |  | GRC |  | ITA |  | MED |  |
| --- | --- | --- | --- | --- | --- | --- | --- | --- |
| | $\sigma$ | $v$ | $\sigma$ | $V$ | $\sigma$ | $v$ | $\sigma$ | $v$ |
| CO<sub>2</sub> | 1.4 | 0.91 | 1.21 | 0.45 | 1.04 | 0.37 | 3.63 | 0.27 |
| CH<sub>4</sub> | 1.9 | 0.76 | 2.3 | 0.54 | 3.34 | 0.45 | 8.72 | 0.34 |
| N<sub>2</sub>O | 0.21 | 1.04 | 0.18 | 0.55 | 0.06 | 0.23 | 0.57 | 0.56 |
| BC | 0.36 | 0.6 | 0.27 | 0.41 | 0.4 | 0.39 | 0.68 | 0.19 |

Emissions for Algeria are highly variable across all species. Looking to the whole Mediterranean domain, BC emissions are the most consistent among the considered inventories, while N_2_O emissions show the highest variability. Our analysis is consistent with recent scientific discussions highlighting the need to align emission inventories to achieve greater accuracy (Cowie et al., 2012; Yona et al., 2020). It is noteworthy that, based on the literature review presented in this work, no emission estimates from atmospheric inversion modelling exist so far for open vegetation fire emissions in the Mediterranean region. Working on the application of this methodology would represent a valuable contribution towards a better quantification (and reduction) of the uncertainties associated with the inventories used to constraint emission estimates in this work.

## 5. Conclusions

This work reviewed the current methodologies for quantifying climate-altering emissions from open vegetation fires in the Mediterranean region, focusing on CO_2_, CH_4_, N_2_O, and BC. Based on our scientific literature review, limited specific efforts have been made to investigate the role of open vegetation fires as a source of climate-altering species in the Mediterranean. The few existing studies were based on a bottom-up approach, while there are no results based on a top-down, observational methodology.

Our review revealed significant deviations in emission estimates from four widely used inventories (GFED v4.1s, GFAS v1.2, FINN v2.5, and EDGAR v8.0), underlying the constraints and limitations on the accounting of climate-altering emissions for this sector. The differences in input data, fire detection, spatial/temporal resolution, and adopted emission factors contribute to the variability in emission quantification between the inventories. FINN v2.5, which uses the satellite data with the higher spatial resolution, is the inventory that provides the highest emission estimates, while the relative differences between the different inventories vary as a function of the species considered, suggesting a role for the different emission factors used. Assuming that the variability among inventories can be considered as a measure of underlying uncertainties in the total emissions, the differences between the inventories suggested a need for further efforts in improving estimates. This is also important when considering anthropogenic fire emissions from the agricultural sector, in particular to determine the relative role of each country’s share in the Mediterranean region.

Inventories like GFED v4.1s, GFAS v1.2 and FINN v2.5 provide high-resolution temporal data, making them valuable for numerical modeling and regional scale assessments. Moreover, with respect to EDGAR (focused on agriculture and waste burning), they provide more comprehensive data about fire emissions, including wildfires. However, their coarse spatial resolution limits their applicability for specific applications like, e.g. city-level modeling. Integrating these inventories with localised datasets, such as high-resolution land-use maps and urban fire observations, is crucial to enhance their utility for city-scale emissions modeling and air quality assessments. Despite these challenges, fire inventories remain indispensable for understanding the atmospheric and climatic impacts of fire emissions.

We hypothesised and discussed the possibility that the inter-annual variability in the emissions can be related to global climate processes, like ENSO. In particular, a possible relationship between the occurrence of La Niña events and the detected emission peaks was discussed. However, the rather short period of time considered in this work, the multiple factors, other than weather conditions that influence the occurrence of forest fires in the Mediterranean, and the complex responses of the extratropical atmosphere to La Niña, require further and more specific studies that go well beyond the objective of this review.

This study highlights the importance of upgrading GHG emission inventories to better reflect the impact of open fire emissions in the Mediterranean region. It highlights significant differences in the reported emissions and underlines the importance of using standardised, transparent, and verifiable methodologies for GHG accounting. The differences between the inventories, especially when down-scaling to the portion of single countries included in the investigated domain suggested further efforts in reducing underlying uncertainties.

In general, we would recommend that future studies should focus on developing consistent methodologies for quantifying and reporting emissions, integrating satellite and ground-based measurement data through atmospheric inversion modelling as already done for other emission sectors. Policy makers should use improved inventories to develop targeted, evidence-based fire prevention and climate mitigation strategies, particularly for the Mediterranean region. This study highlights the need for accurate emissions data and its integration into policy frameworks to support effective climate change mitigation at regional and local levels.

## Acknowledgments

We would like to thank the teams behind the GFED v4.1s, GFAS v1.2, FINN v2.5, and EDGAR v8.0 inventories for providing the crucial data that significantly contributed to our research. ERA5 data were provided by ECMWF through Copernicus Climate Change Service (C3S). We gratefully thank Annalisa Cherchi (CNR - ISAC) for the useful discussion about the role of ENSO in the Mediterranean region. We thank Riccardo Testolin (ISAFOM-CNR and NBFC, Italy) for providing the Mediterranean mask. We acknowledge the support from the University School for Advanced Studies IUSS Pavia (“Scuola Universitaria Superiore IUSS Pavia”) and the Institute of Atmospheric Sciences and Climate (ISAC), National Research Council (CNR), Italy. We also acknowledge the project funded by the European Union – NextGenerationEU under the National Recovery and Resilience Plan (NRRP), Mission 4 Component 2 Investment 1.4 - Call for tender No. 3138 of December 16, 2021, rectified by Decree n.3175 of December 18, 2021 of the Italian Ministry of University and Research under award Number: Project code CN_00000033, Concession Decree No. 1034 of June 17, 2022 adopted by the Italian Ministry of University and Research, CUP B83C22002930006, Project title “National Biodiversity Future Centre - NBFC”. During this research work, Rabia Ali Hundal and Saurabh Annadate were supported by the Italian national inter-university PhD course in Sustainable Development and Climate change (PhD SDC, link: www.phd-sdc.it).

## Declaration of AI-assisted technologies in the writing process

During the preparation of this work the authors used DeepL (https://www.deepl.com/) in order to check the English language. After using this tool, the authors reviewed and edited the content as needed and take full responsibility for the content of the publication.

